# Enhancer heterogeneity in acute lymphoblastic leukemia drives differential gene expression between patients

**DOI:** 10.1101/2024.12.08.627394

**Authors:** Alastair L. Smith, Nicholas Denny, Catherine Chahrour, Kim Sharp, Natalina Elliott, Joe Harman, Thomas Jackson, Huimin Geng, Owen Smith, Jonathan Bond, Irene Roberts, Ronald W. Stam, Nicholas T Crump, James O.J. Davies, Anindita Roy, Thomas A. Milne

## Abstract

Genetic alterations alone cannot account for the diverse phenotypes of cancer cells. Even cancers with the same driver mutation show significant transcriptional heterogeneity and varied responses to therapy. However, the mechanisms underpinning this heterogeneity remain under-explored. Here, we find that novel enhancer usage is a common feature in acute lymphoblastic leukemia (ALL). In particular, *KMT2A::AFF1* ALL, an aggressive leukemia with a poor prognosis and a low mutational burden, exhibits substantial transcriptional heterogeneity between individuals. Using single cell multiome analysis and extensive chromatin profiling, we reveal that much transcriptional heterogeneity in *KMT2A::AFF1* ALL is driven by novel enhancer usage. Using high resolution Micro-Capture-C in primary patient samples, we also identify patient-specific enhancer activity at key oncogenes such as *MEIS1* and *RUNX2*, driving high levels of expression of both oncogenes in a patient-specific manner. Overall, our data show that enhancer heterogeneity is highly prevalent in *KMT2A::AFF1* ALL and may also be a mechanism that drives transcriptional heterogeneity in cancer more generally.

**Key Points:**

- Leukemia patients with the same driver mutations often display gene expression differences
- Using chromatin profiling and high resolution 3C methods we show that enhancer heterogeneity drives gene expression differences

## Introduction

Although genetic alterations drive carcinogenesis, they alone cannot account for the diverse phenotypes of cancer cells ^1^. Even cancers that contain the same driver mutation can display significant transcriptional heterogeneity and varied responses to therapy ^2–5^. However, the mechanisms underpinning transcriptional heterogeneity remain under-explored.

In higher eukaryotes, gene transcription initiates from promoters ^6^, but highly regulated context-specific expression is often controlled by the activity of distal regulatory elements termed enhancers ^6,7^. Multiple models have been proposed for enhancer function (reviewed in ^7^), but key attributes of enhancer activity include a specific chromatin profile (enrichment for H3K27ac and H3K4me1); the production of short bidirectional transcripts termed eRNAs; the binding of sequence-specific transcription factors; the tendency to come within close proximity of the promoter in 3D space when active; and the ability to drive tissue- and temporal-specific gene expression ^6–10^.

Aberrant enhancer activity is being increasingly recognized as a driver of human disease, the most obvious examples involving large genomic rearrangements that bring enhancers close to the genes they aberrantly regulate ^11,12^. Recent work has also identified cancer-specific regulatory programs driven by tumor-associated enhancers ^13,14^. DNA insertions and single-nucleotide polymorphisms (SNPs) can alter transcription factor (TF) binding and thus impact enhancer function ^11,13^, but less is known about how non-DNA mutations such as epigenetic changes can drive cancer-specific enhancer function.

Acute lymphoblastic leukemia (ALL) is the most common type of childhood cancer ^15,16^. Although children with ALL generally respond well to therapy, there are specific subtypes that still confer a poor prognosis, especially in infant ALL ^16,17^, although new therapies such as Blinatumomab show great promise frontline treatment for infant ALL ^18^. Even so, there are limited treatment options for ALL patients who relapse ^19^ and even when successful, treatment of childhood ALL can have life-long adverse impacts ^20,21^. ALL caused by rearrangements of the *KMT2A* gene (formerly *MLL*) producing in-frame gene fusions that create novel fusion proteins ^22^. The most common *KMT2A* rearrangement (*KMT2A*r) is *KMT2A::AFF1* ^22^ which is the major cause of infant ALL and where it has a near silent mutational landscape ^17,23–26^. *KMT2A* rearrangements also cause acute myeloid leukemias (AML) as well as ALL in older children and adults ^22^. Although not as mutationally silent as infants, even in older patients, *KMT2A* rearrangements have a much lower mutational burden than other subtypes of ALL or AML and lack commonly recurring mutations ^25,27,28^.

Despite the low mutational burden, *KMT2A*r leukemias exhibit substantial transcriptional and phenotypic heterogeneity between individuals ^4,25^, providing an ideal model to study transcriptional heterogeneity without the confounding effect of numerous cooperating mutations. KMT2A fusion proteins are generally thought to drive disease through both transcriptional and epigenetic mechanisms, primarily by binding to the promoters of genes ^29,30^. We have recently shown that the KMT2A::AFF1 protein complex drives high levels of gene expression by inducing oncogenic enhancer activity ^31,32^. However, it is unknown how much enhancer usage might differ between patients of the same leukemia subtype, or how differential enhancer usage might drive individual transcription patterns and thereby influence prognostic outcomes.

Here, in B-ALL primary patient samples and cell lines with a variety of molecular subtypes, including *KMT2A::AFF1*, *ETV6::RUNX1*, *DUX4*/*ERG* and hyperdiploidy, we identified extensive transcriptional heterogeneity and, by extensive chromatin profiling, revealed the existence of a large number of novel enhancers in individual patients. CRISPR-Cas9-based deletion of representative enhancers unique to individual KMT2A::AFF1 cell lines altered target gene expression in a cell-specific manner. For the first time, we were able to use the high resolution 3C technique Micro-Capture-C (MCC ^33^) in a primary patient sample to show in detail that KMT2A::AFF1-bound enhancers near *MEIS1* and *RUNX2* contact the *MEIS1* and *RUNX2* promoters, directly implicating these patient-specific enhancers in the overexpression of these genes. Taken together, our data suggest that enhancer heterogeneity is highly prevalent in *KMT2A::AFF1* ALL and that this likely plays a significant role in the phenotypic diversity observed between patients in ALL.

## Methods

### Patient samples

Infant (<1 year old at diagnosis) and childhood (1–18 years old) ALL samples were obtained from Blood Cancer UK Childhood Leukaemia Cell Bank (now VIVO Biobank, UK) under their ethics approval (REC: 23/EM/0130), and from Our Lady’s Children’s Hospital, Crumlin, Ireland (REC: 21/LO/0195). Informed consent was obtained from all participants or those with parental responsibility.

### ATAC-seq and TOPmentation

ATAC-seq was conducted on 5 × 10^4^ live cells using Nextera Tn5 transposase (Illumina) as previously described ^31^. Libraries were sequenced by paired-end sequencing with a 75-cycle high-output Nextseq 500 kit (Illumina). TOPmentation was performed as described ^31^. See supplemental Materials for protocol and data analysis details.

### Single Cell Multiome

Cryopreserved bone marrow cells were thawed, and 16,000 live CD19+ blasts of 4 *KMT2A- AFF1+* ALL patient samples were FACS-sorted and nuclei were extracted using the recommended protocol (Chromium). Male/female sample pairs were loaded together on one Chromium 10x lane for processing with the Single Cell Multiome ATAC + Gene Expression protocol. Sequencing was performed on a NovaSeq 6000, with gene expression libraries sequenced on a PE150 S4 flow cell and ATAC-seq libraries on an SP PE50 flow cell. See supplemental Materials for data analysis.

### RNA-seq and qRT-PCR

For qPCR, RNA was extracted from 1×10^6^ cells with the RNeasy Mini Kit (Qiagen). Reverse transcription was conducted using Superscript III (ThermoFisher Scientific) with random hexamer primers (ThermoFisher Scientific), and cDNA was analyzed by Taqman qPCR, using the housekeeping gene *YWHAZ* for gene expression normalization.

### Micro-Capture-C

Micro-Capture-C was performed on patient sample chALL1 as described ^34^ (also see supplemental Materials). Analysis was performed using the MCC pipeline ^33^.

### Whole genome sequencing (WGS)

Genomic DNA was extracted from 5×10^6^ cells using a Monarch Genomic DNA extraction kit (NEB). 2 ng of genomic DNA was incubated at 55°C for 15 minutes with 0.4 ul TDE1 (Illumina) before purification (Qiagen MinElute PCR purification kit). Indexing and sample purification was performed in the same manner as for TOPmentation. Libraries were sequenced on a NovaSeq X (2×150 bp). Data analysis details are provided in supplemental Materials.

## Results

### Altered enhancer activity regulates differential gene expression in KMT2A::AFF1 ALL cells

To investigate how altered enhancer activity regulates gene expression, we first focused on a specific B-ALL subtype, KMT2A::AFF1-driven leukemia, which is characterized by a minimal mutational burden^25^. To explore the mechanistic aspects of differential enhancer activity in greater detail, we compared gene expression differences in two *KMT2A::AFF1* ALL patient-derived cell lines, SEM and RS4;11. RNA sequencing of these cell lines revealed 4,351 differentially expressed genes (Figure 1A), indicating significant transcriptional heterogeneity and validating them as suitable models for further study.

**Figure 1.**
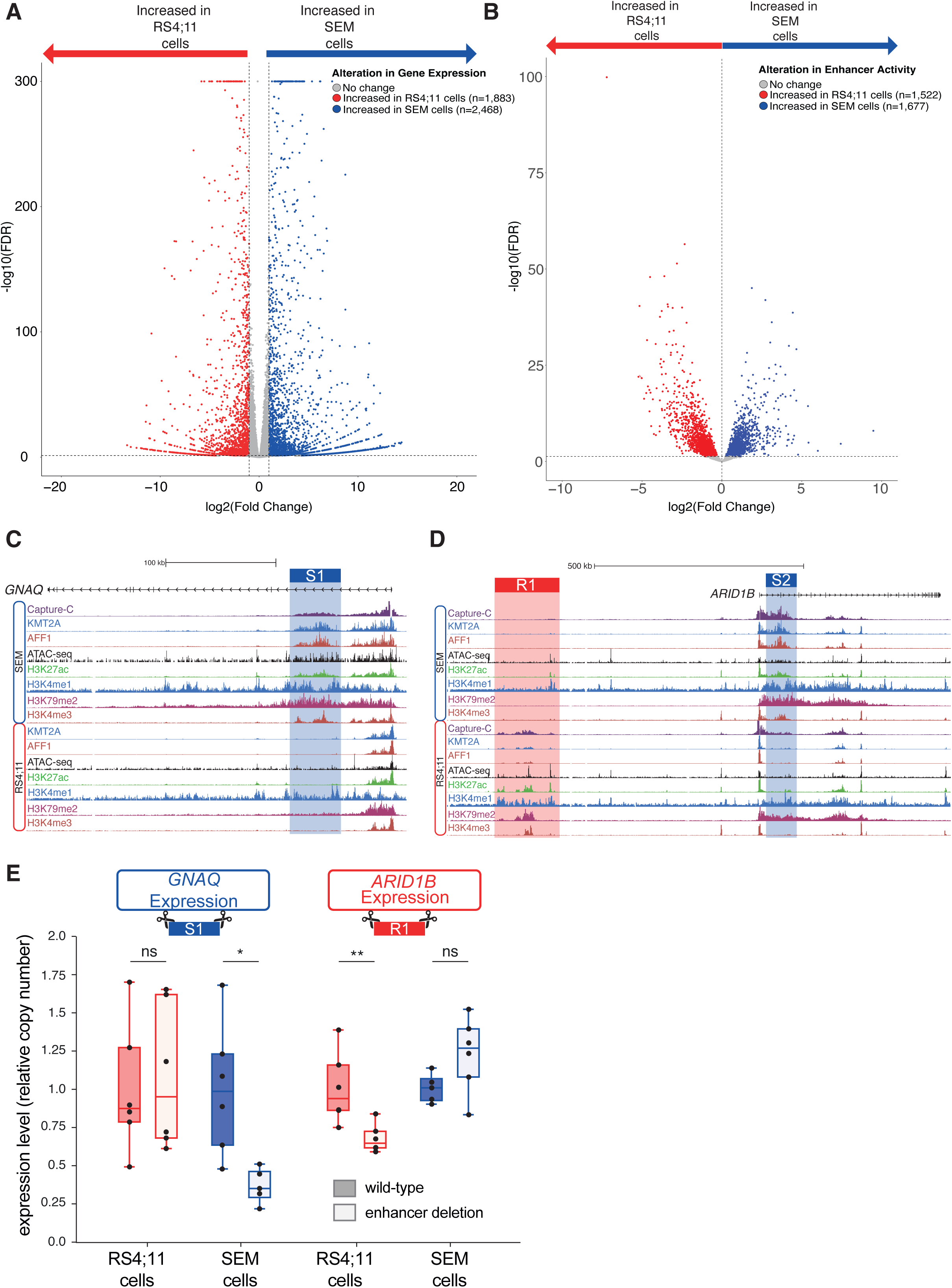
Differential enhancer regions in *KMT2A::AFF1* cell lines are functional enhancers that drive differential gene expression. (A) Volcano plot of differentially expressed genes between RS4;11 (1,883; red) and SEM cells(2,468; blue) or no significant change (gray) from three biological replicates, FDR <0.05. (B) Volcano plot of enhancers with significantly increased accessibility in RS4;11 (1,522; red) or SEM cells (1,677; blue) or enhancers with unaltered accessibility (gray) from eight biological replicates, FDR <0.05. (C) ChIP-seq tracks at the *GNAQ* locus for KMT2A, AFF1, H3K27ac, H3K4me1, H3K79me2 and H3K4me3 together with ATAC-seq and Capture-C in SEM cells using the *GNAQ* promoter as a viewpoint. The SEM specific *GNAQ* enhancer (S1) is highlighted in blue. (D) ChIP-seq tracks at the *ARID1B* locus for KMT2A, AFF1, H3K27ac, H3K4me1, H3K79me2 and H3K4me3 together with ATAC-seq and Capture-C in SEM cells using the *ARID1B* promoter as a viewpoint. The RS4;11 specific intergenic enhancer (R1) is highlighted in red, and the SEM specific intragenic enhancer (S2) is highlighted in blue. (E) RT-qPCR comparing the expression of enhancer deletion mutants (non-filled) to wild type (filled) in RS4;11 (red) or SEM cells (blue) when deleting either the intragenic GNAQ enhancer (S1; left) or ARID1B intergenic enhancer (R1). Significance of alterations in relative copy number were determined by a two-sided T-test with correction for multiple testing (Benjamini-Hochberg), n=6 biological replicates. * adjusted p-value <0.05, ** <0.01

To compare enhancer usage between the two cell lines, we measured chromatin accessibility, a proxy for enhancer activity, under the assumption that increased accessibility correlates with transcription factor binding. Even though both cell lines are derived from *KMT2A::AFF1* leukemias, significant differences in enhancer accessibility were seen between SEM and RS4;11 cells (Figure 1B). After filtering, 7,194 putative enhancers were identified and of these 1,677 displayed significantly increased accessibility in SEM cells, with 1,522 increased in RS4;11 cells (Figure 1B). Although the majority of the differential enhancers displayed a relatively minor difference in accessibility, with 925 and 1124 enhancers displaying less than a two-fold increase in RS4;11 and SEM cells, respectively, a significant number of enhancers were uniquely detected in one of these cell lines.

Strikingly, we identified enhancers with differential activity at known KMT2A::AFF1 target genes such as *GNAQ* and *ARID1B* ^31^. The *GNAQ* gene is associated with intragenic enhancers in SEM cells which are absent in RS4;11 cells (Figure 1C, blue shading). In contrast, at *ARID1B*, an upstream intergenic enhancer is present in RS4;11 cells but absent in SEM (Figure 1D, red shading), and an intragenic enhancer in SEM cells shows reduced activity in RS4;11 (Figure 1D, blue shading). At both of these genes, the cell line-specific enhancers show an increased frequency of interaction with the promoter, as measured by Capture-C (Figure 1C-D), a typical feature of enhancers^10^.

We validated the activity of the differential *GNAQ* and *ARID1B* enhancers by targeting them for CRISPR-Cas9-mediated deletion (Supplemental Figure 1A). Deletion of the *GNAQ* intragenic enhancer locus in SEM cells decreased expression of *GNAQ* in SEM cells, while in RS4;11 cells no significant change in *GNAQ* expression was observed (Figure 1E). Similarly, deletion of the RS4;11-specific *ARID1B* enhancer sequence in RS4;11 cells resulted in a reduction in gene expression, whereas RNA levels were not significantly altered in SEM deletion mutants (Figure 1E). This is consistent with these regions being active enhancers in a cell line-specific manner.

To explore the effect of cell line-specific enhancers on transcription genome-wide, we linked enhancers to the nearest gene. Enhancers displaying increased activity were more frequently linked to genes displaying increased expression (51.8% of enhancer-gene pairs in SEM cells and 64.9% in RS4;11 cells, Supplemental Figure 1B), arguing that these epigenetic differences are functionally relevant. In addition, genes linked to one or more differentially active enhancer(s) displayed significantly increased differences in gene expression (Supplemental Figure 1C). Thus, these regions are *bona fide* enhancers with differential effects on gene expression.

### Heterogeneity in enhancer activity is common in B-ALL

We next wanted to determine if the enhancer heterogeneity we observed in the cell line models also applied in ALL patient samples. First, we leveraged publicly available RNA-seq and ATAC-seq datasets from a diverse cohort of 24 primary patient samples^35^ comprised of *ETV6::RUNX1*, *DUX4*/*ERG* and hyperdiploid B-ALL subtypes. We examined the variance in gene expression both between and within the B-cell ALL subtypes and observed substantial transcriptional variability (Supplemental Figure 2A). While each of the three subtypes clustered separately as expected, using either hierarchical or k-means clustering we identified six distinct clusters of genes based on their expression profile, implying substantial variability within the patient subtypes. For example, expression patterns in clusters 1 and 6 divided patient samples in the hyperdiploid subtype into two distinct subtypes (Supplemental Figure 2A).

To establish the source of this transcriptional heterogeneity we examined the chromatin accessibility landscape of these samples. We generated a consensus peak set consisting of 71,800 open chromatin regions and categorised these open chromatin sites into promoter (<2.5 kb from the nearest transcription start site (TSS)) and putative enhancers (≥ 2.5 kb; these will also include non-enhancer loci) and correlated normalized read counts between samples at both promoter and enhancer regions Importantly, promoter peaks exhibited a substantially higher correlation (0.81-0.96; Figure 2A) between samples compared to enhancers (0.52-0.91; Figure 2B), supporting the proposition that variability in enhancer usage could distinguish leukemia subtypes. To further establish this, enhancer regions were able to separate the B-ALL subtypes into their distinct subsets, whereas promoter regions failed to fully distinguish the *ETV6::RUNX1* and *DUX4*/*ERG* subtypes. This shows that enhancer activity might provide a more robust signature of leukemia subtype (Figure 2C), consistent with enhancers being the main source of transcriptional differences ^36–38^.

**Figure 2.**
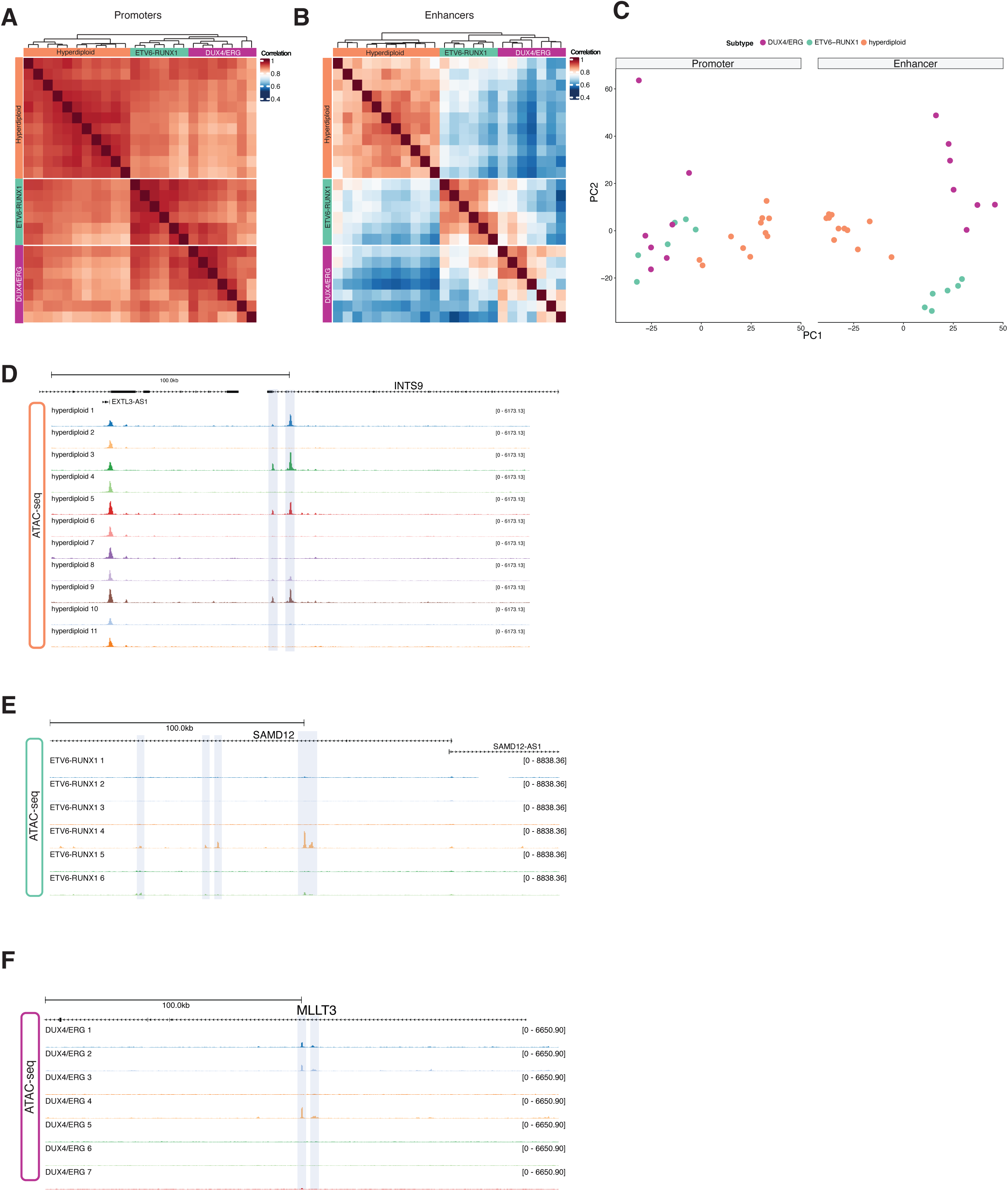
B-ALL patients display enhancer heterogeneity between individuals. (A) Correlation of accessibility at promoter regions (<2.5 kb from a TSS) as measured by ATAC-seq signal between DUX4/ERG ETV6-RUNX1 and hyperdiploid subtypes. (B) Correlation of accessibility at putative enhancers (> 2.5 Kb from a TSS) as measured by ATAC-seq signal between DUX4/ERG ETV6-RUNX1 and hyperdiploid subtypes. (C) PC analysis of chromatin accessibility at promoters (left) and enhancers (right) for all three B-ALL subgroups. (D) ATAC-seq at the INTS9 locus for hyperdiploid samples. Putative enhancer regions with a high degree of inter-sample variability are highlighted in blue. (E) ATAC-seq at the *MLLT3* locus for DUX4/ERG samples. Putative enhancer regions with a high degree of inter-sample variability are highlighted in blue. (F) ATAC-seq at the *SAMD12* locus for ETV6-RUNX1 samples. Putative enhancer regions with a high degree of inter-sample variability are highlighted in blue.

We next wanted to refine this analysis and determine whether there was variability in enhancer usage between individual patients. To do this, we intersected the open chromatin regions identified within each sample to the consensus peak set (Supplementary Figure 2B). Across all subtypes, promoter regions displayed a higher degree of concordance with 45.6-61.2% being accessible in all samples within each subtype. In contrast, putative enhancer regions exhibited a substantially decreased consistency between samples with only 11.9-28.9% of regions being accessible in all samples, implying a high degree of variability in the activity of enhancers between samples. Examples of the variability in enhancer activity could be readily observed at the *INTS9* locus in the hyperdiploid subtype (Figure 2D), the *SAMD12* locus in the *ETV6::RUNX1* subtype (Figure 2E) and the *MLLT3* locus in the *DUX4*/*ERG* subtype (Figure 2F). Taken together, these results show two things: 1) different leukemia subtypes share a distinct enhancer signature, and 2) despite this shared enhancer signature, there is also significant variability between samples, suggesting patient to patient heterogeneity.

We next aimed to determine if these changes in enhancer activity could be linked directly to gene expression. We compared scaled chromatin accessibility at these enhancers with scaled gene expression of the nearest gene for each patient. Despite the limitation that enhancers do not always regulate the nearest promoter, we observed a notable positive correlation (ranging from 0.36 to 0.62) between putative enhancer accessibility and gene expression for the most variable genes (n=100, Supplementary Figure 2C). This correlation suggests that differential enhancer activity likely contributes to the transcriptional heterogeneity observed between these patient samples.

### There is extensive enhancer heterogeneity in *KMT2A::AFF1* B-ALL patient leukemia samples

To identify heterogenous enhancer usage in *KMT2A::AFF1* patient blasts we wanted to be able to compare enhancer usage with gene expression on a single cell level. To do this, we performed 10x single cell (sc) Multiome (ATAC + RNAseq) on FACS-sorted CD19+ blast populations obtained from four *KMT2A::AFF1* ALL patient samples (3 infants and 1 older child). Using the scATAC modality (Figure 3A), we examined the degree of heterogeneity in chromatin accessibility and identified 6,231 regions of open chromatin with significantly altered accessibility between leukemic blasts from the 4 patients (340,494 regions identified in total; Figure 3B). The majority (64%) of differentially accessible regions were located more than 2.5 kb from the nearest TSS (Figure 3C), predominantly consisting of intronic elements (Figure 3D), implying that most of these regions are putative enhancers.

**Figure 3.**
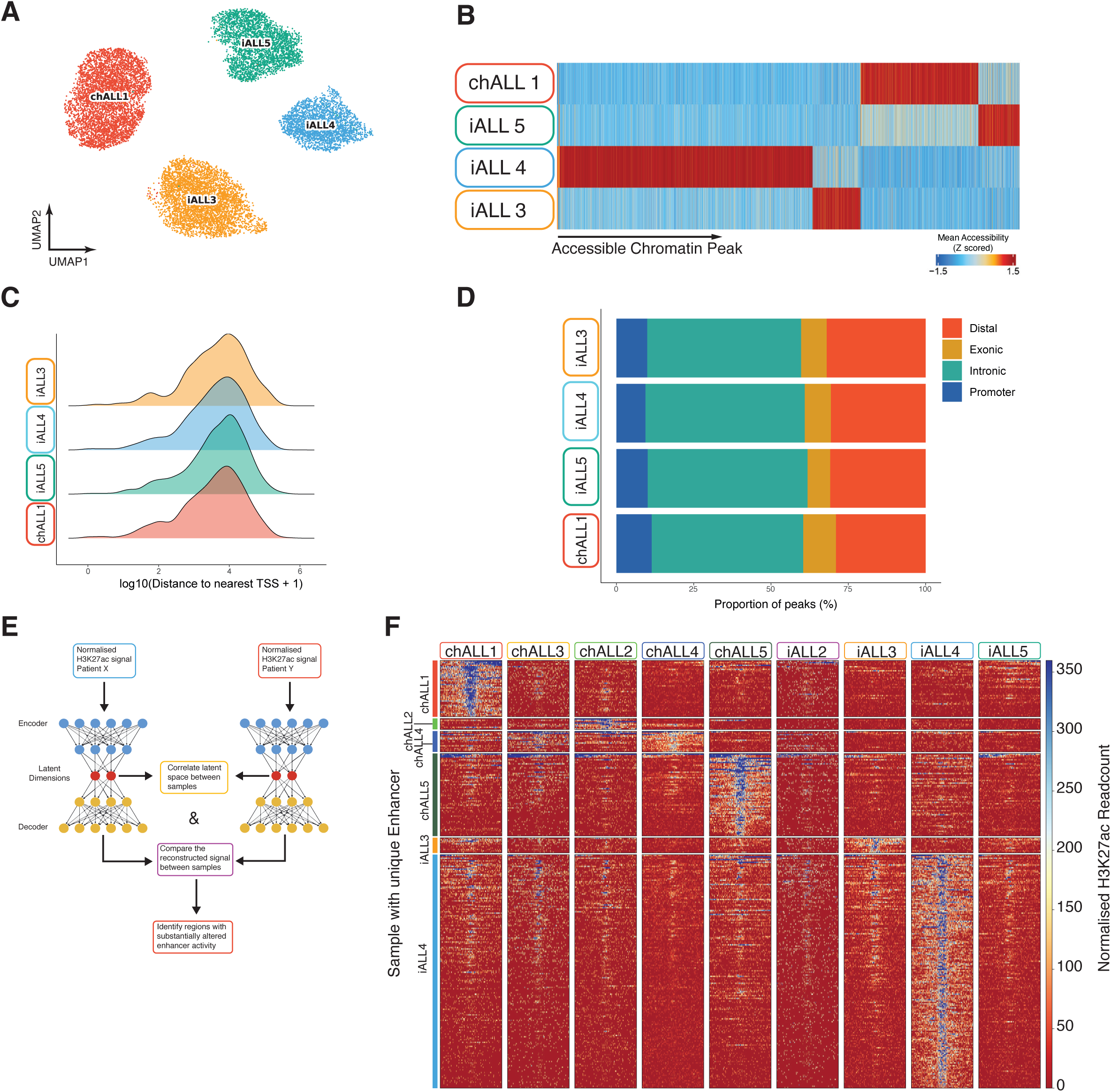
Differential enhancer regions in *KMT2A::AFF1* patients are readily observed. (A) UMAP of the scATAC-seq modality for four KMT2A::AFF1 blast samples (B) ATAC-seq peaks (accessible chromatin) displaying significantly increased accessibility in one out of the four patient samples. (C) Genomic distribution of uniquely accessible ATAC-seq peaks relative to the nearest TSS (transcription start site). (D) Annotation of the genomic location of unique ATAC-seq peaks (E) Schematic of the strategy used to identify unique enhancer peaks using H3K27ac ChIP-seq datasets. (F) Tornado plot of H3K27ac signal in KMT2A::AFF1 patient samples at enhancers identified as being patient-specific.

To robustly identify active enhancers, we performed detailed epigenetic profiling in nine *KMT2A::AFF1* patient samples (four infants and five children) with a low-cell number optimized chromatin immunoprecipitation protocol, TOPmentation^31^ (Supplemental Table 1), including the 4 patients with sc-multiome data. We defined a consensus set of putative enhancer regions by identifying H3K27ac enriched regions that were ≥2.5kb from the nearest TSS. Due to the limited material available, resulting in a lack of replicates and variability in sample viability, we devised an autoencoder-based strategy to limit sample-sample noise and identify enhancer regions that displayed substantially altered enhancer activity, marked by increased levels of H3K27ac, between patient blasts (Figure 3E). This allowed us to identify 290 patient-specific putative enhancers across the nine patient samples (Figure 3F; Supplemental Figure 3A-F). In many cases, these patient-specific enhancers were associated with the binding of KMT2A (Supplemental Figure 3A-F).

Since we observed patient-specific enhancer activity, we sought to determine the extent of transcriptional heterogeneity between *KMT2A::AFF1* patient samples. To this end, we integrated our single nucleus gene expression data with a published single cell gene expression (scGEX ^4^) dataset of *KMT2A::AFF1* blasts from three iALL patients to increase the number of patients in our dataset.. Initial dimensionality reduction, following batch correction (Figure 4A), indicated that each patient sample formed a distinct cluster, implying unique gene expression profiles. Moreover, analysis of differential gene expression between the samples (Figure 4B; FDR < 0.01) revealed a surprising number of genes (3,306) that exhibited heterogeneous expression, with top marker genes shown in Figure 4B. This number is comparable to the differential expression observed between *KMT2A::AFF1* cell lines (Figure 1A). Interestingly, marker gene analysis indicated that two key *KMT2A::AFF1* target genes, *MEIS1* and *RUNX2*, were both highly elevated in chALL1 (Figure 4B-C, Supplemental Figure 4A). Returning to our epigenetic data from this patient, we observed a chALL1 unique putative enhancer region downstream of *MEIS1* (Figure 4D) and a putative enhancer upstream of *RUNX2* in chALL1 and iALL2 (Figure 4E), both of which were enriched for KMT2A binding.

**Figure 4.**
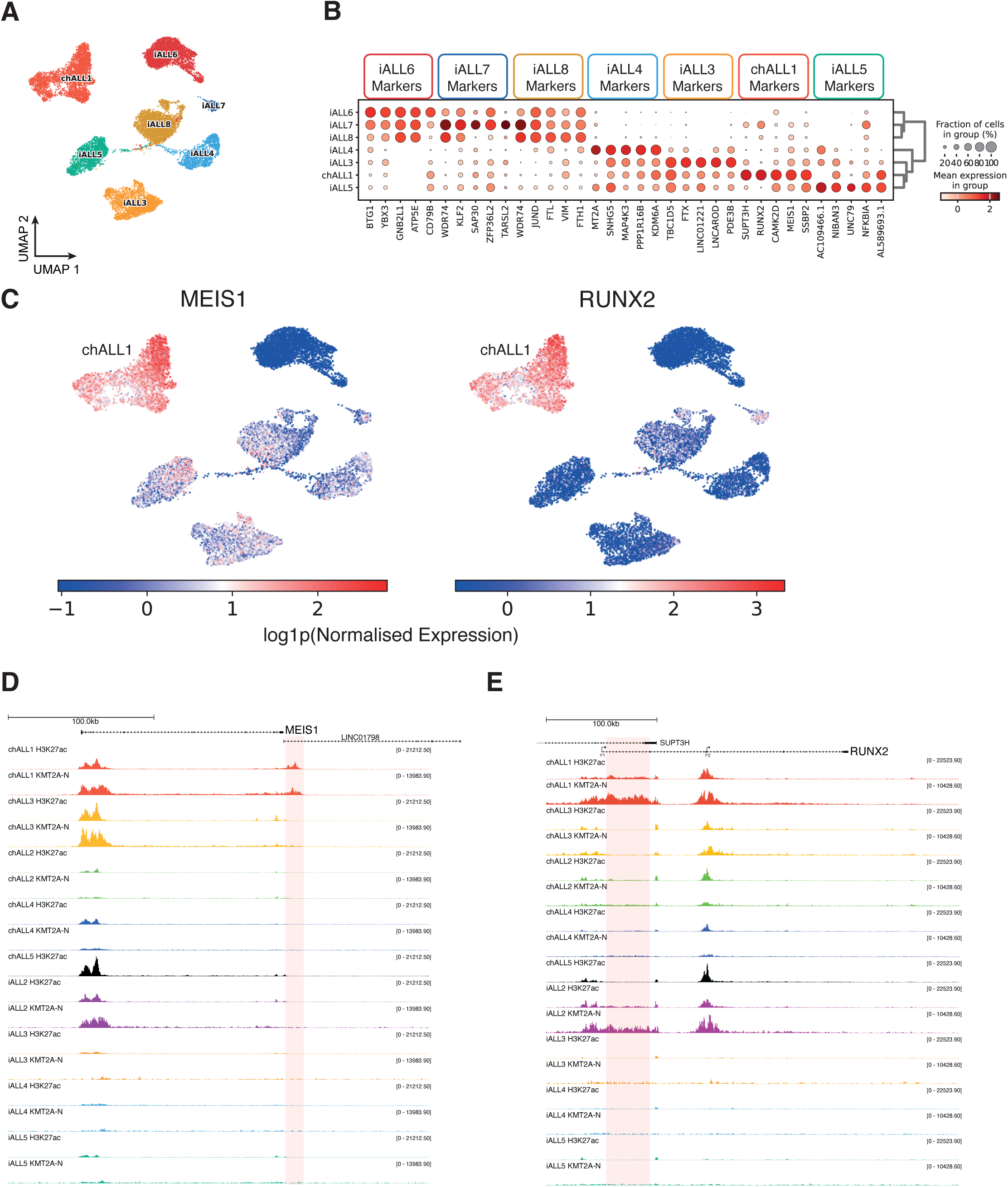
Differential enhancer activity in *KMT2A::AFF1* patients drives oncogene specific expression such as at *MEIS1* and *RUNX2*. (A) UMAP of snGEX for four KMT2A::AFF1 blast samples (chALL1, iALL3-5) and three scGEX samples. (B) Dotplot of marker gene analysis between seven KMT2A::AFF1 blast samples, showing the top five marker genes per sample. (C) Normalised MEIS1 (left) and RUNX2 (right) expression in KM2TA::AFF1 sn/scGEX samples. (D) TOPmentation for H3K27ac and the N-terminus of KMT2A (KMT2A-N) in KMT2A::AFF1 blast samples at the *MEIS1* locus. The chALL1 unique enhancer region downstream of *MEIS1* is highlighted in red. (E) TOPmentation for H3K27ac and KMT2A-N in KMT2A::AFF1 blast samples at the *RUNX2* locus. The chALL1 and iALL2 specific enhancer region upstream of *RUNX2* is highlighted in red.

Both *MEIS1* and *RUNX2* expression are increased upon relapse in B-ALL (Supplemental Figure 4B-C and ^39^), and expression of *RUNX2* is significantly higher in patients that did not exhibit complete remission (Supplemental Figure 4D). High *MEIS1* expression is also significantly correlated with a worse outcome in childhood ALL (Supplemental Figure 4E), while high *RUNX2* expression significantly correlates with a worse overall survival in *non- KMT2Ar* adult ALL (Supplemental Figure 4F). Together, these results show that high levels of expression of two key genes associated either with relapse or worse overall survival (i.e. *RUNX2* and *MEIS1*), can be driven by enhancer activity that is specific to an individual patient leukemia sample.

### Novel enhancers in *KMT2A::AFF1* leukemias are directly linked to genes that they regulate

Our analysis identified multiple unique enhancers in patient samples. To confirm these regions were *bona fide* enhancers, we performed the high resolution 3C method Micro-Capture-C (MCC^33^), using 75 promoters as viewpoints, in chALL1 blasts, in which we identified putative unique enhancers at *MEIS1* and *RUNX2*. This revealed that these enhancer elements frequently came into physical proximity with the promoters of *MEIS1* and *RUNX2* (Figure 5A-B) strongly implying that the enhancer elements regulate these genes. We did not observe promoter interactions with the chALL1 *MEIS1* enhancer in the SEM cell line, confirming that these interactions are dependent on the presence of the active enhancer (Figure 5A). Interestingly, we do observe the presence of the *RUNX2* enhancer in SEM cells (Figure 5B) in addition to iALL2 (Figure 4E), implying that this enhancer is not strictly chALL1 specific but instead is variable among KMT2A::AFF1 patients.

**Figure 5.**
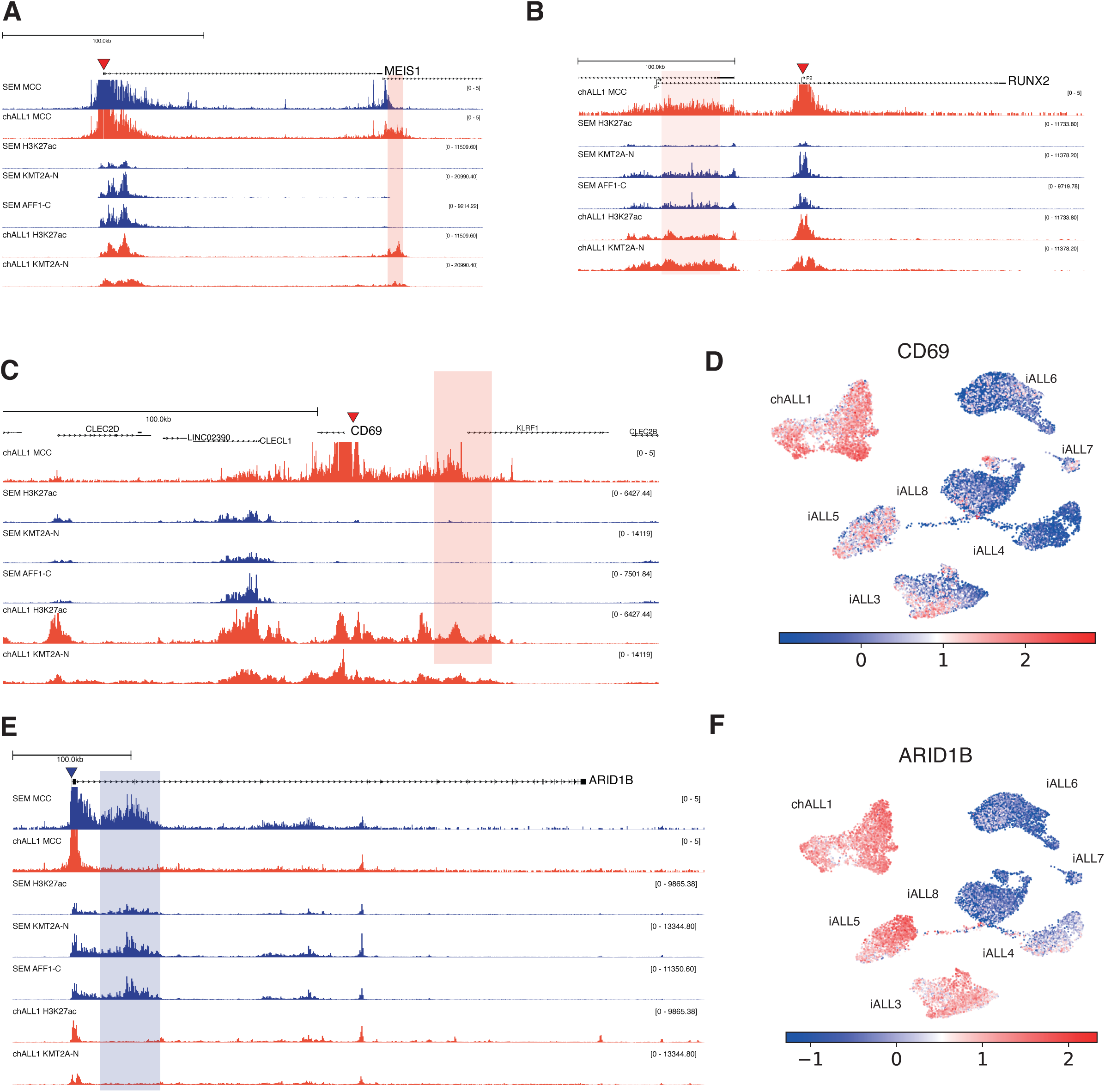
MCC reveals patient-specific enhancer-promoter interactions in primary patient cells. (A) MCC for the *MEIS1* viewpoint (red triangle) in chALL1 and SEM cells together with TOPmentation for H3K27ac and KMT2A-N at the *MEIS1* locus. The chALL1 specific enhancer downstream of *MEIS1* is highlighted in red. (B) MCC for the *RUNX2* viewpoint (red triangle) together TOPmentation for H3K27ac and KMT2A-N at the *RUNX2* locus. The enhancer present in chALL1 upstream of the RUNX2 promoter is highlighted in red. (C) MCC for the *CD69* viewpoint (red triangle) together with ChIP-seq for H3K27ac and KMT2A-N at the *CD69* locus. The chALL1 specific enhancer is highlighted in red. (D) Expression of *CD69* in KMT2A::AFF1 patient blast samples. (E) MCC for the *ARID1B* viewpoint (blue triangle) together with ChIP-seq for H3K27ac and KMT2A-N at the *ARID1B* locus. The SEM specific intragenic enhancer is highlighted in red. (F) Expression of *ARID1B* in KMT2A::AFF1 patient blast samples.

Using our MCC data we were able to confirm the targets of additional chALL1 unique enhancers, e.g. at the *CD69* locus (Figure 5C). In agreement with our hypothesis that these patient-specific enhancers increase the expression of their target gene, *CD69* expression was elevated in chALL1 (Figure 5D, Supplemental Figure 5A). Interestingly, patient specific enhancers have previously been identified at the *CD69* gene in cases of mixed phenotype acute leukemia^40^. In contrast to *MEIS1*, *RUNX2* and *CD69*, *ARID1B* expression was not substantially increased in chALL1 when compared to other patient samples, likely because an *ARID1B* intragenic enhancer that is present in SEM cells is missing in chALL1 (Figure 1D, Figure 5E,F, Supplemental Figure 5B). Taken together, our data confirm that the differential enhancer regions identified at *MEIS1*, *RUNX2* and *CD69* are indeed enhancers for these genes, and thus implicating them in oncogene overexpression.

### KMT2A::AFF1 binding is predictive of enhancer activity

Having identified differential enhancer usage in *KMT2A::AFF1* leukemias, we wanted to confirm that it was not driven by small-scale changes in DNA sequence at these enhancers. Using WGS we looked for single nucleotide variants (SNVs) within the enhancers exhibiting differential activity in the KMT2A::AFF1 cell lines (SEM and RS4;11). Only 25-43.2% of enhancers contained at least one SNV or insertion-deletion (INDEL) and frequencies were comparable between common and cell line-unique enhancers (Figure 6A), implying that mutations in enhancer regions are unlikely to explain most of the differential enhancer activity. Furthermore, we examined if any of these variants exhibited allele-specific bias in our ATAC-seq data, as increased chromatin accessibility might indicate a functional consequence of the mutation. Strikingly, only 0.5-2.2% of enhancers contained one or more heterozygous SNVs exhibiting allele-specific bias (Figure 6B). Taken together, these data suggest that SNVs in these enhancer regions are unable to explain the differences in enhancer activity observed.

**Figure 6.**
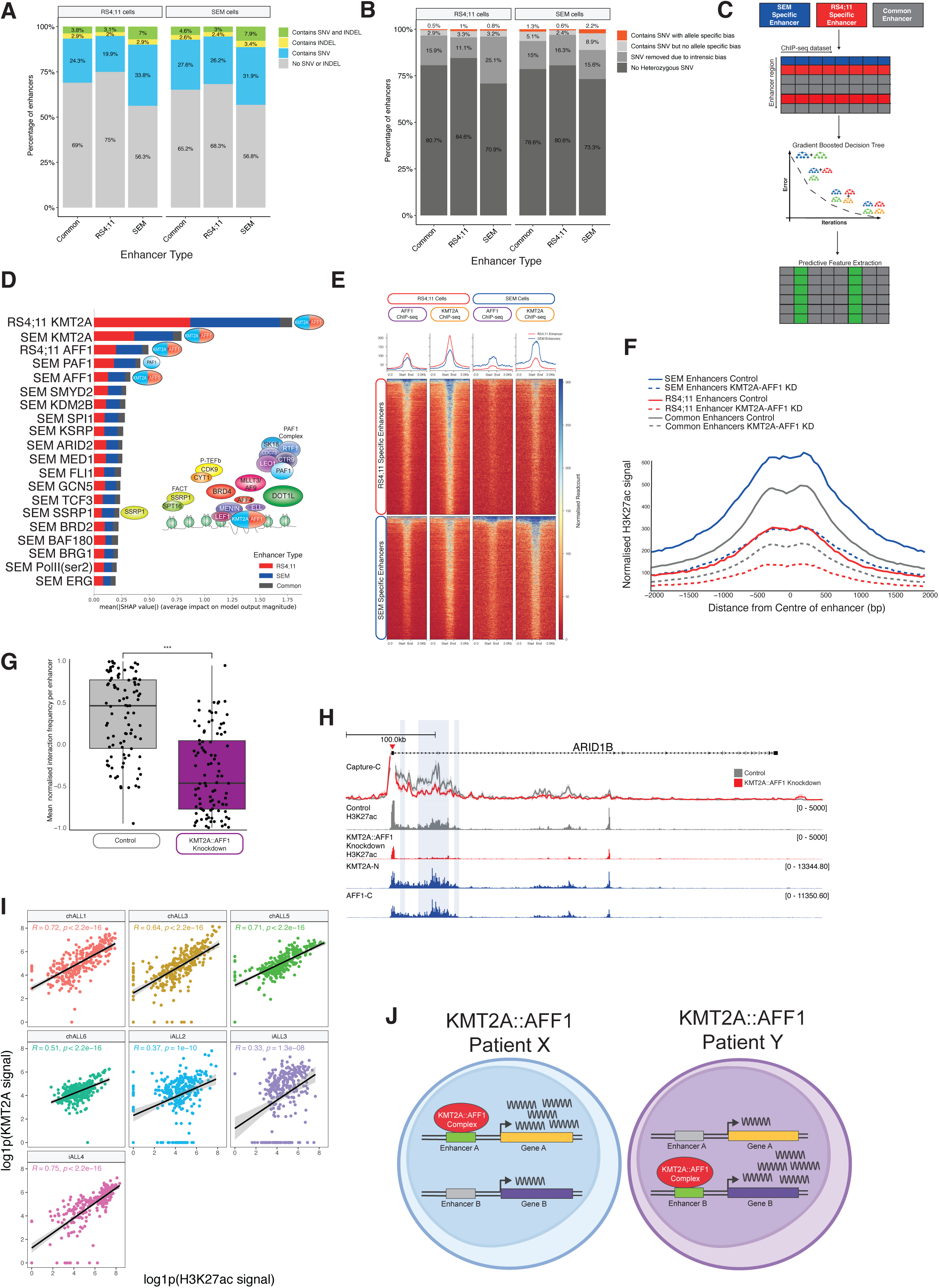
An unbiased machine learning model identifies KMT2A::AFF1 complex binding as a driver of differential enhancer usage. (A) Proportion of enhancers (Common = no change in activity, RS4;11 = increased activity in RS4;11 cells, SEM = increased activity in SEM cells) containing either a SNV (blue), INDEL (yellow) or both (green) in RS4;11 cells (left) or SEM cells (right). (B) Proportion of enhancers of each enhancer type (Common, RS4;11, SEM) containing no heterozygous SNVs (dark grey), SNVs removed due to intrinsic bias (e.g. problematic genomic regions or mapping bias; lighter grey), SNVs without allele specific bias in accessibility (light grey) or those exhibiting allele specific bias in accessibility as measured by ATAC-seq (red). (C) Schematic of the strategy used to determine key predictive features of differential enhancer activity. ChIP-seq signal for 56 factors was extracted over enhancers with increased activity in RS4;11 cells (red; 1,522), or SEM cells (blue; 1,677) or those common to both (gray; 4232). A gradient boosted decision tree was trained from this data and predictive features were extracted using SHAP. (D) The relative feature importance for each enhancer category (increased activity in SEM cells (blue), RS4;11 cells (red) or common enhancers (gray)) of the top 20 most important feature for differential enhancer prediction. Features that correspond to binding of the KMT2A::AFF1 complex are highlighted and a schematic of the complex is provided for reference. (E) Tornado plot of AFF1 and KMT2A ChIP-seq signal at enhancers displaying increased activity in RS4;11 cells (top) or SEM cells (bottom) in RS4;11 (left) or SEM (right) cells. (F) H3K27ac ChIP-seq signal at enhancers with increased activity in SEM cells (blue), RS4;11 cells (red) or common enhancers (gray) upon KMT2A::AFF1 knockdown by siRNA (dashed line). (G) Change in enhancer-promoter interaction frequency of enhancers with increased activity in SEM cells following siRNA based KMT2A::AFF1 knockdown assessed by Capture-C. Interaction frequency for each enhancer-promoter pair is shown relative to the mean interaction frequency of the control; n=3 biological replicates per condition. (H) Example of a loss of enhancer- promoter interactions at the *ARID1B* locus as assessed by Capture-C in control (grey) or KMT2A::AFF1 knockdown conditions (red) in three biological replicates. Enhancers with increased activity in SEM cells are highlighted in blue. ChIP-seq for H3K27ac in control (grey) or KMT2A::AFF1 knockdown conditions (red) in addition to the N-terminus of KMT2A and the C-terminus of AFF1 are provided for reference. (I) Pearson correlation between H3K27ac and KMT2A signal at the 290 blast-specific enhancers identified for each patient sample. (J) Model for the role of the KMT2A::AFF1 complex in promoting transcription heterogeneity between patients.

Our past work has shown that KMT2A::AFF1 binding is essential for maintaining enhancer function at a shared set of KMT2A::AFF1 target genes ^31^. The unique enhancers identified in this current study often have KMT2A bound to them (using an antibody that also detects KMT2A::AFF1), implicating KMT2A::AFF1 in driving enhancer heterogeneity. To explore this in an unbiased manner, we created a model to identify the most likely factors driving enhancer heterogeneity in KMT2A::AFF1 ALL. We developed a gradient-boosted decision tree model using 56 lab-generated ChIP-seq datasets from SEM and RS4;11 cells (Figure 6C, Supplementary Table 2) to be able to predict which enhancers would exhibit increased activity in SEM or RS4;11 cells (weighted F1 score of 0.845 for predicting enhancers increased in RS4;11 and 0.736 SEM specific enhancers; Supplementary Figure 6A-B), as well as those with unchanged activity. We used this model to identify the datasets with the highest predictive power (Figure 6D) and verified that the five most predictive features displayed significant enrichment differences between common and cell type-specific enhancer classes (Supplementary Figure 6C).

Strikingly, the model identified the presence of KMT2A in each cell line as the most important feature for defining differential enhancer usage (Figure 6D) followed by its fusion partner AFF1 and the KMT2A::AFF1 complex component PAF1 ^31,41,42^. Taken together this implicates binding of the KMT2A::AFF1 complex with differential enhancer usage. In line with this, KMT2A and AFF1 ChIP-seq signal in both cell lines was enriched at cell-specific active enhancers and depleted at inactive enhancers (Figure 6E). This is also consistent with our observation of KMT2A binding at patient-specific enhancers (Supplementary Figure 3A-F)

Finally, to establish a causative role for KMT2A::AFF1 at these differential enhancers, we next examined the effect of siRNA mediated knockdown of KMT2A::AFF1 in SEM cells ^31^ on the level of H3K27ac at these elements, as a proxy for enhancer activity (Figure 6F). We observed a substantial reduction in the level of H3K27ac at all enhancer subtypes (common, SEM-specific and RS4;11-specific) upon KMT2A::AFF1 knockdown. This reduction was matched by a decrease in enhancer-promoter interaction frequency by Capture-C (Figure 6G,H, Supplementary Figure 6D), with 75.23% of enhancers within 500 kb of the 35 promoters analysed exhibiting a significantly reduced interaction frequency (Figure 6G). Furthermore, returning to our primary sample specific enhancers, we observed a positive correlation between H3K27ac signal intensity and KMT2A binding at the unique enhancers in each patient (R=0.33-0.75; Figure 6I) in further support of our model predictions. Taken together, these data implicate the binding of KMT2A::AFF1 in the activity of these cell line- and patient-specific enhancers, suggesting that differential binding of the KMT2A::AFF1 complex is a key driver of transcriptional heterogeneity due to the regulation of differential enhancer activity (Figure 6J).

## Discussion

Much work has gone into developing targeted therapies that are effective but relatively non-toxic. Unfortunately, many patients still either fail to respond or relapse for reasons that are not always clear. Cancers with the same driver mutations show varied responses to therapy, along with significant transcriptional heterogeneity. Here, we have shown that even a relatively mutationally silent cancer subtype such as *KMT2A::AFF1* can display extensive transcriptional heterogeneity, driven in part by the usage of patient-specific leukemia enhancers. When important oncogenes such as *MEIS1* and *RUNX2* are overexpressed due to novel enhancer usage, this may have implications for therapeutic response and patient outcomes in leukemia. Variability in enhancer activity has been observed across various cancers, including gastric adenocarcinoma ^43^, prostate tumours ^44^, and luminal breast cancer ^45,46^, suggesting that this may represent a broader mechanism contributing to heterogeneity across cancer types.

A key question arising from our findings is the origin of these novel enhancers. One hypothesis is that patient-specific SNPs may drive differential enhancer activity by altering transcription factor binding sites, a mechanism proposed for donor-derived lymphoblastoid lines ^47^. However, our analysis of sequencing data from the SEM and RS4;11 cell lines revealed only a small number of SNPs associated with novel enhancers, despite the extensive epigenetic differences observed. This suggests that genetic variation alone is unlikely to fully account for the emergence of patient-specific enhancers in this context.

Another possibility is that the target cell in which the leukemia originates may provide unique and distinct epigenetic landscapes for KMT2A::AFF1 binding, and thus create the possibility of differential enhancer usage. Since each *KMT2A::AFF1* ALL patient displayed a distinct pattern of enhancer usage, cell type is unlikely to fully explain the effects observed, unless there are multiple possible cell types of origin, one for each patient. Instead, it is likely that a stochastic element contributes to initial enhancer activation, and is subsequently reinforced by stabilization of KMT2A::AFF1 binding. For example, KMT2A::AFF1 binding has been shown to be altered by DNA methylation ^39,48^, so stochastic and developmental stage-specific differences in DNA methylation patterns in progenitor cells could be a driver of differential binding and enhancer usage.

A deeper exploration of the source of these unique enhancers will require more extensive investigation, but this work establishes variable enhancer usage as a common driver of transcriptional heterogeneity in multiple leukemias, suggesting this could be one common mechanism underpinning aspects of patient heterogeneity across different cancer subtypes.

## Acknowledgements

T.A.M. and A.L.S, were funded by Medical Research Council (MRC, UK) Molecular Haematology Unit grant MC_UU_00016/6 and MC_UU_00029/6. C.C. is funded by a Wellcome Trust Genome Medicine and Statistics studentship award. This project was further supported by the Fight Kids Cancer Funding Programme, supported by Imagine For Margo, Fondation KickCancer, Foundatioun Kriibskrank Kanner, Federazione Italiana Associazioni Genitori e Guariti Oncoematologia Pediatrica and Cris Cancer Foundation. J.R.H. was funded by an Engineering & Physical Sciences Research Council (EPSRC) Doctoral Training Program grant project, reference 2119788 and EP/R513295/1. N.T.C. was supported by a Kay Kendall Leukaemia Fund Intermediate Fellowship (KKL1443). Primary haematological malignancy samples used in this study were provided by Blood Cancer UK Childhood Leukemia Cell Bank. J.O.J.D. is supported by the Lister Institute, the MRC Molecular Haematology Unit (MC_UU_00029/04), Wellcome (225220/Z/22/Z) and the Oxford National Institute of Health Research Biomedical Research Centre (NIHR203311).N.D. is funded by Cancer Research UK (SEBCATP-2022/100011). AR is supported by a Wellcome Trust Clinical Research Career Development Fellowship (216632/Z/19/Z) and Medical Research Council (MRC, UK) Molecular Haematology Unit grant MC_UU_00029/7.

## Authorship Contributions

A.L.S., N.T.C. and T.A.M. conceived the experimental design. A.L.S., N.D., K.S., N.E., J.H. and T.J. carried out experiments. A.L.S., C.C., J.H., T.J., H.G. and N.T.C. analyzed and curated the data. A.L.S., N.D., A.R. and T.A.M. interpreted the data and wrote the manuscript. O.S., J.B., R.W.S. and A.R. provided resources. I.R., J.O.J.D., A.R. and T.A.M. provided funding and supervision. A.L.S., N.D., C.C., J.H., H.G., O.S., J.B., I.R., R.W.S., N.T.C., J.O.J.D., A.R., T.A.M., reviewed and edited the manuscript.

## Supplementary Material

### Supplemental Figure Legends

**Supplemental Figure 1 –.**
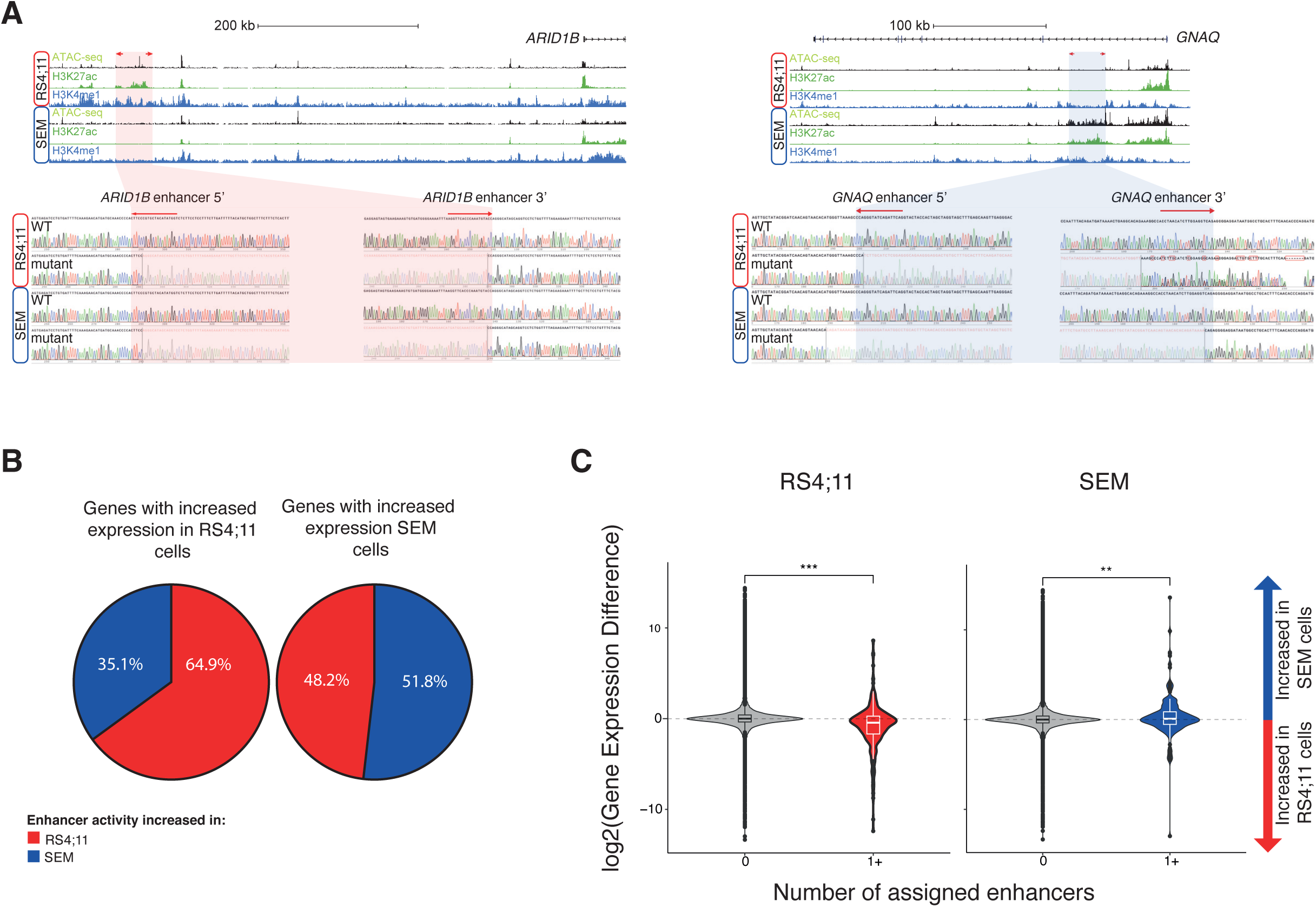
(A) ATAC-seq and ChIP-seq for H3K4me1 and H3K27ac in RS4;11 and SEM cells at the ARID1B and GNAQ loci. The enhancers that were deleted are highlighted in red and the locations of sgRNAs are shown with red arrows. Sanger sequencing results of the mutant enhancers are shown below. (B) Proportion of genes displaying increased expression in RS4;11 (left) and SEM (right) cells, associated with enhancers displaying increased activity in RS4;11 (red) or SEM (blue) cells. (C) Comparison of the gene expression difference between RS4;11 (left) and SEM (right) cells for genes that are associated with one or more enhancers displaying increased activity in RS4;11 cells (red) or SEM cells (blue). Positive values indicate increased gene expression in SEM cells and negative indicate increased expression in RS4;11 cells. *** p< 0.001.

**Supplemental Figure 2 –.**
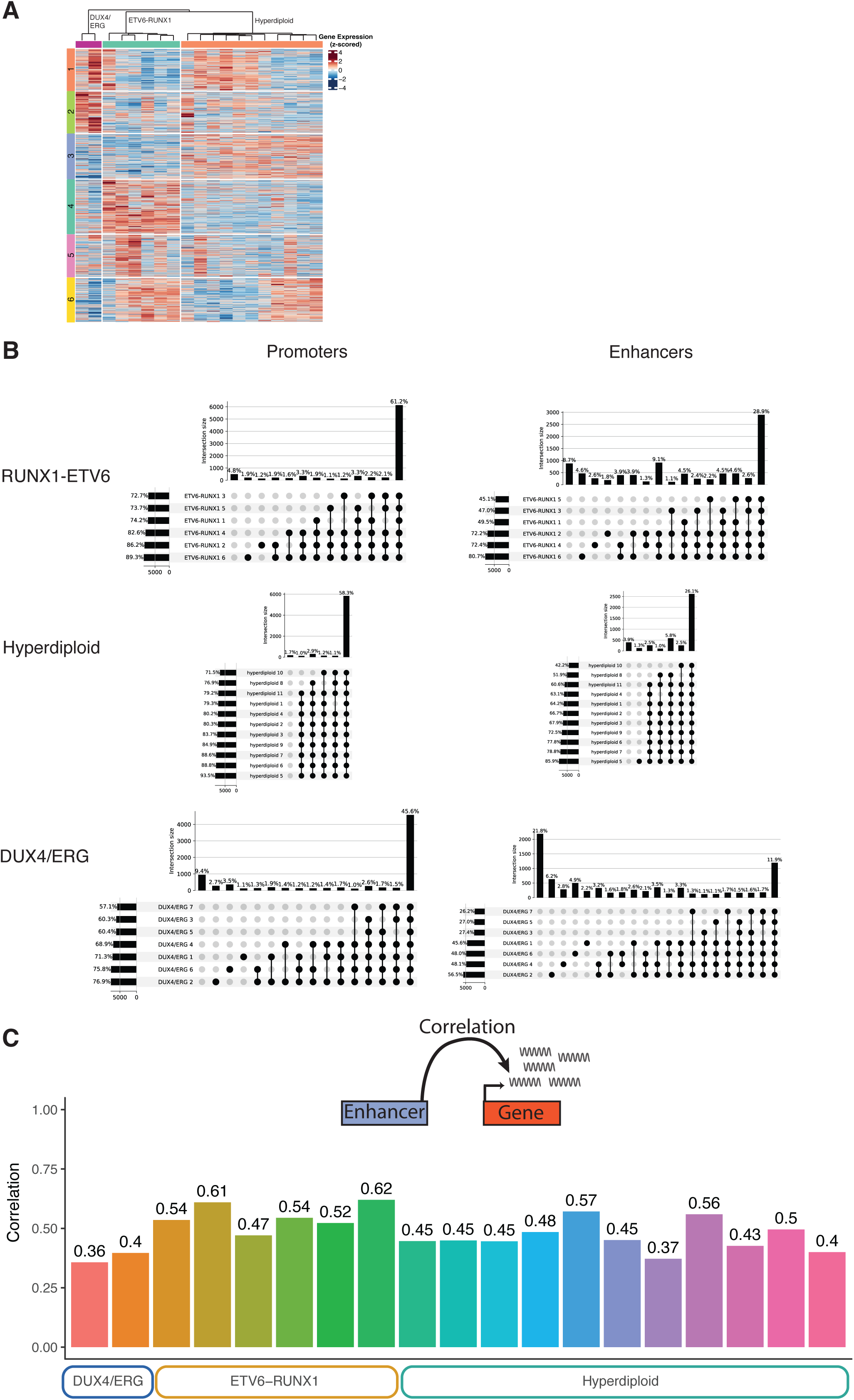
(A) Normalised gene expression of three B-ALL subtypes (DUX4/ERG - green, ETV6-RUNX1 - red and hyperdiploid - blue). Genes have been clustered using K-means. (B) Upset plots for accessible chromatin regions at promoters and putative enhancer regions for the three B-ALL subgroups. (C) Correlation between gene expression and chromatin accessibility for the 100 genes with the highest degree of variability within the subgroups.

**Supplemental Figure 3 –.**
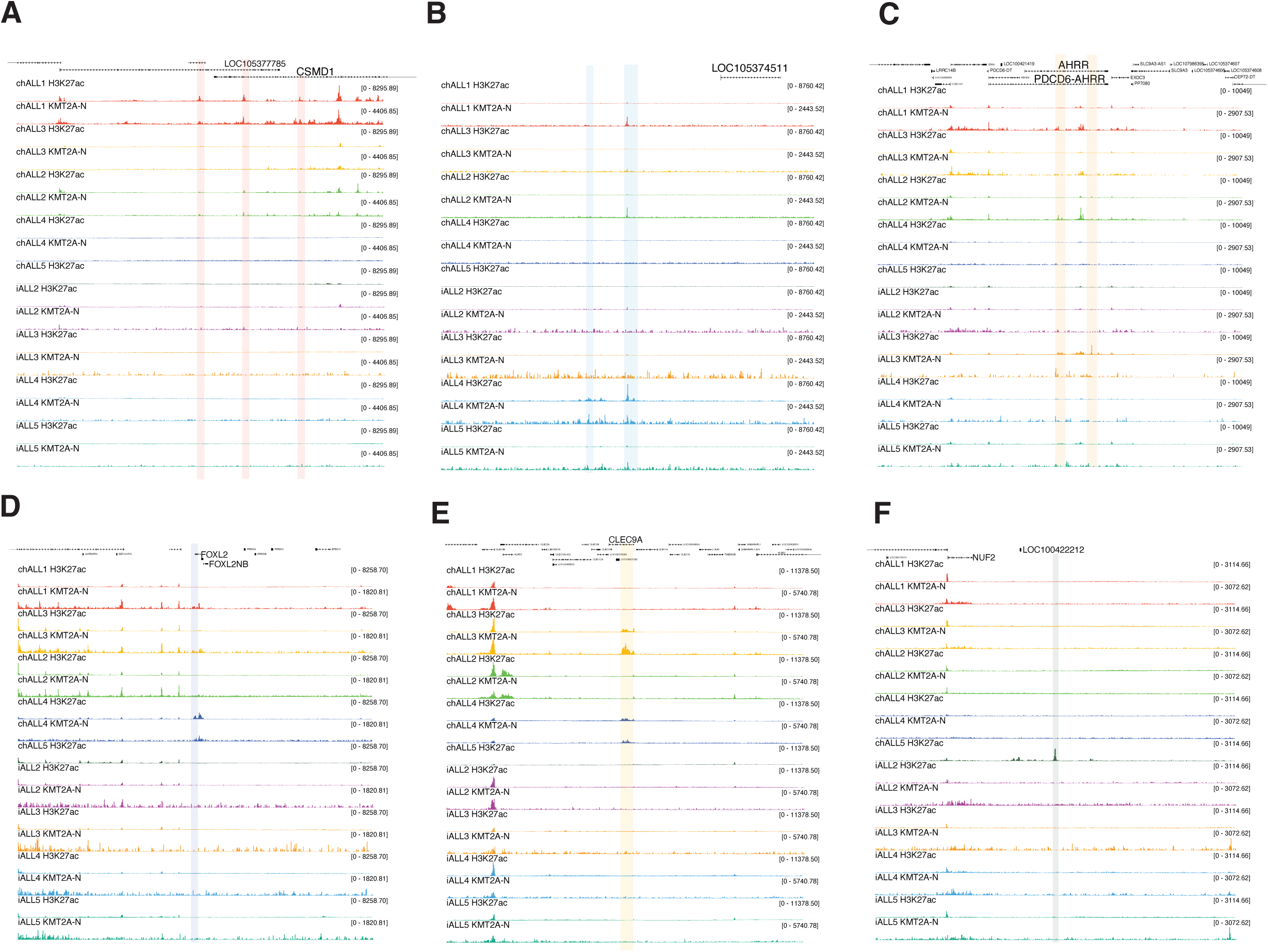
(A) TOPmentation for H3K27ac and KMT2A at the CSMD1 locus. Identified chALL1 specific enhancers are highlighted in red. (B) TOPmentation for H3K27ac and KMT2A. Identified iALL4 specific enhancers are highlighted in blue. (C) TOPmentation for H3K27ac and KMT2A at the AHRR locus. Identified iALL3 specific enhancers are highlighted in orange. (D) TOPmentation for H3K27ac and KMT2A at the FOXL2 locus. Identified chALL4 specific enhancers are highlighted in blue. (E) TOPmentation for H3K27ac and KMT2A at the CLEC9A locus. Identified chALL3 specific enhancers are highlighted in yellow. (F) TOPmentation for H3K27ac and KMT2A upstream of the NUF2 locus. Identified chALL5 specific enhancer is highlighted in grey.

**Supplemental Figure 4 –.**
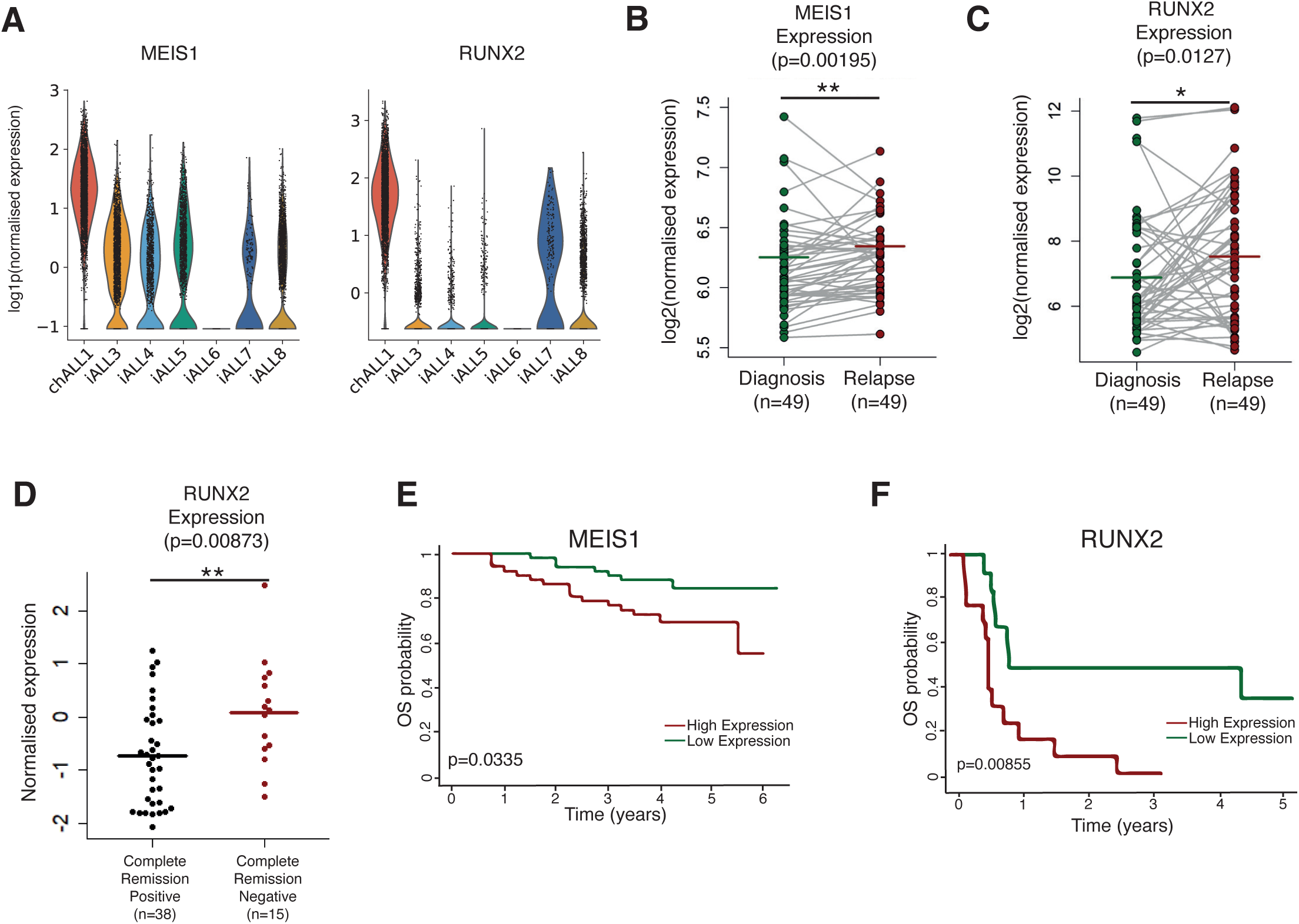
(A) MEIS1 (left) and RUNX2 (right) Gene expression in KMT2A::AFF1 patient blasts. (B) MEIS1 expression at paired diagnosis and relapse. ** p<0.01. Data from COG P9906 n=49 pairs of relapse vs diagnosis B-ALL (GSE28460). P-values were calculated using a two-sided paired Wilcoxon test. (C) RUNX2 expression at paired diagnosis and relapse. * p<0.05. Data from COG P9906 n=49 pairs of relapse vs diagnosis B-ALL (GSE28460). P-values were calculated using a two-sided paired Wilcoxon test. (D) RUNX2 expression for B-ALL patients that experienced complete remission (left) compared to those that did not (right). ** p<0.01. Data from ECOG E2993 adult B-ALL clinical trial (GSE5314, n=54). P-values were calculated using a two-sided Wilcoxon test. (E) Survival curve comparing high (red) and low (green) MEIS1 expression in B-ALL. Data analyzed from COG P9906 childhood B-ALL clinical trial. (F) Survival curve comparing high (red) and low (green) RUNX2 expression in B-ALL. Data analyzed from ECOG E2993 adult B-ALL clinical trial.

**Supplemental Figure 5 –.**
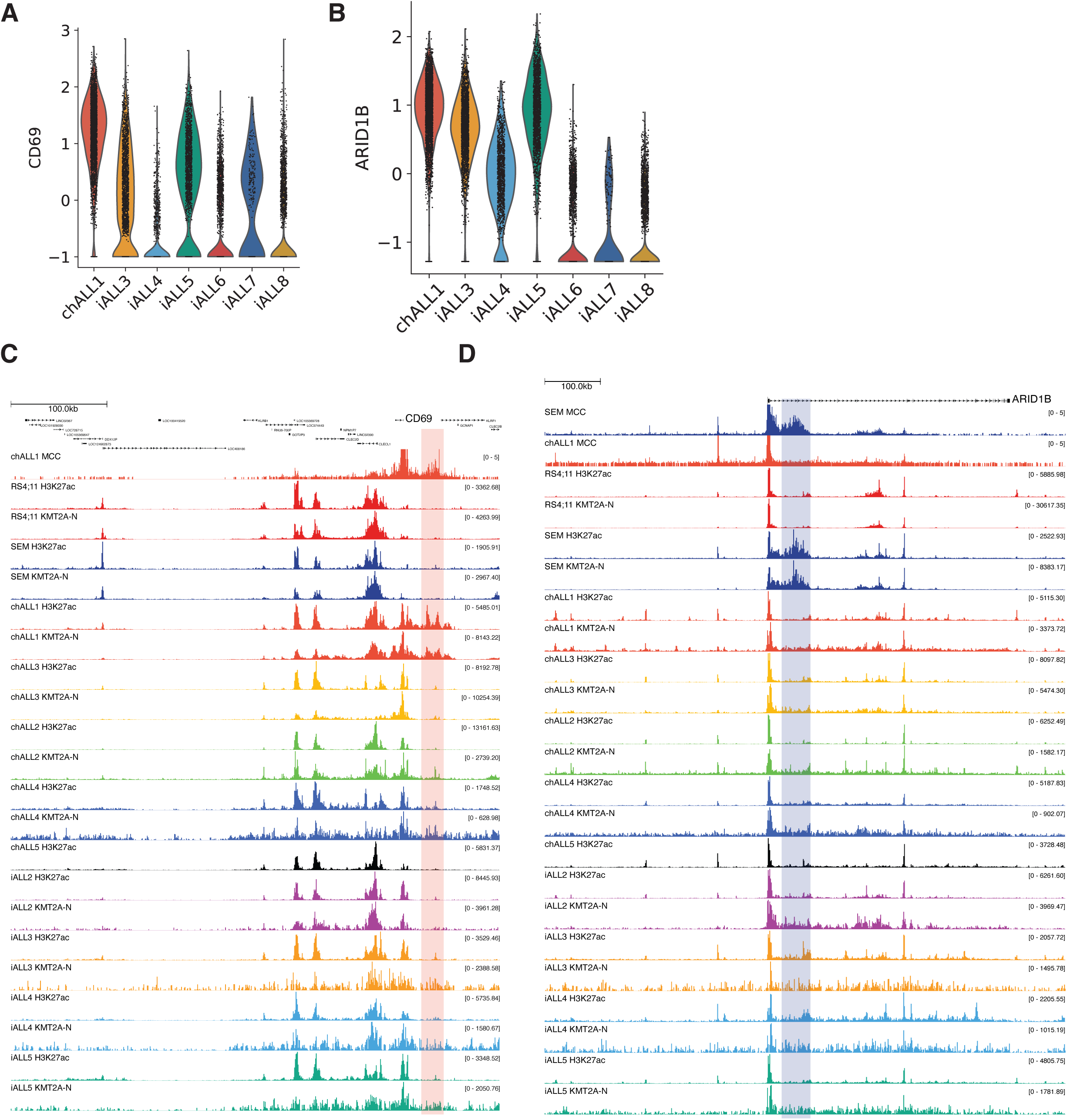
(A) CD69 expression in KMT2A::AFF1 patient blasts. (B) ARID1B expression in KMT2A::AFF1 patient blasts (C) MCC in chALL1 and ChIP-seq (SEM and RS4;11 cells) together with TOPmentation for H3K27ac and the N-terminus of KMT2A for SEM, RS4;11 and other patient blast samples at the CD69 locus. The chALL1 specific enhancer region is highlighted in red. (D) MCC in SEM cells and in chALL1 together with ChIP-seq (SEM and RS4;11 cells) and TOPmentation for H3K27ac and the N-terminus of KMT2A for SEM, RS4;11 and other patient blast samples at the ARID1B locus. The SEM specific intragenic enhancer region is highlighted in blue.

**Supplemental Figure 6 –.**
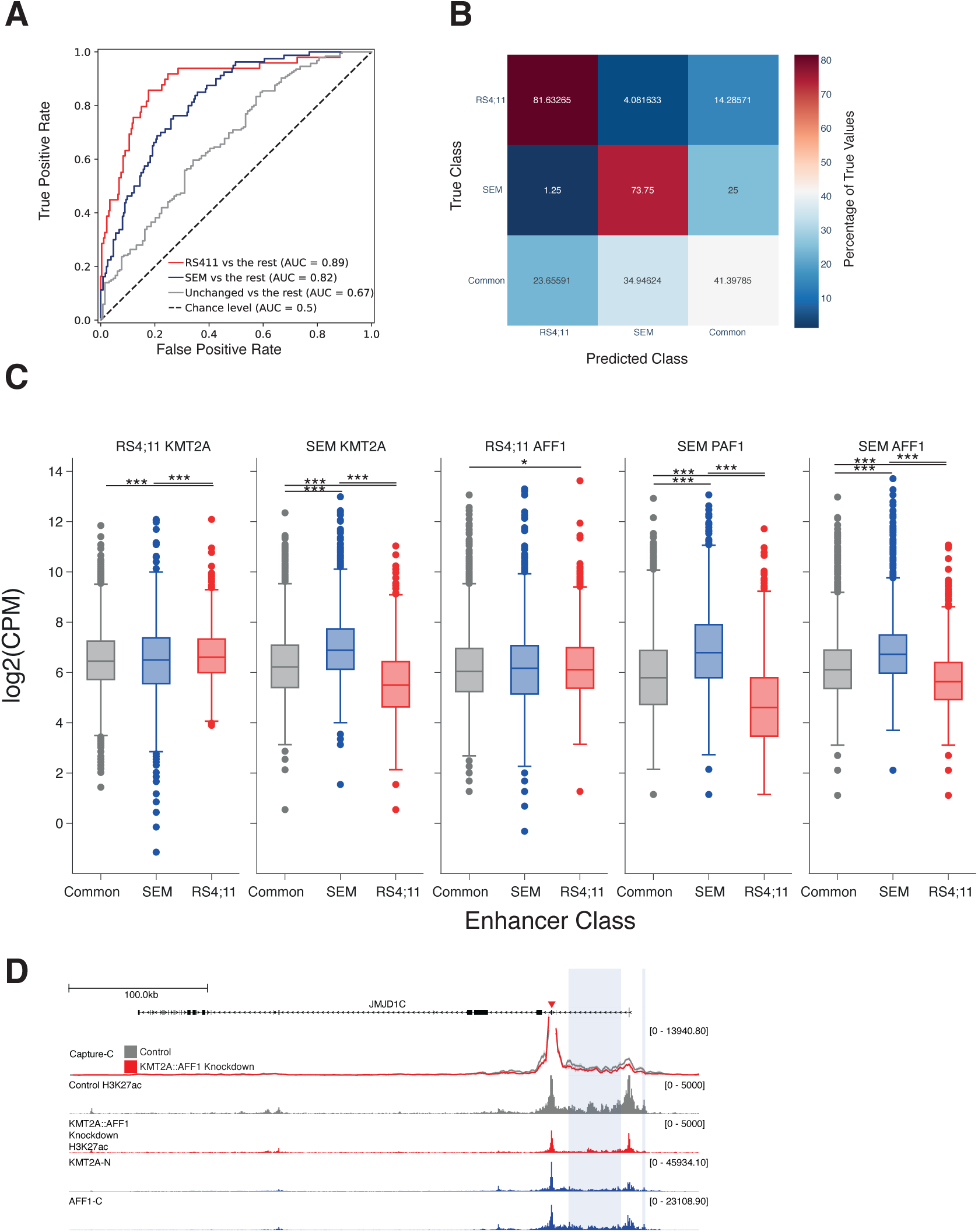
(A) Receiver operating characteristic (ROC) curves from the differential enhancer prediction model. Area under the curve (AUC) scores are provided for each classification. (B) Confusion matrix of enhancer classification predictions from held-out test data, normalised to the number of true positive examples. (C) Box plots for the top five most important features for enhancer classification. * p < 0.05, ** p <0.01, *** p <0.001¬ (D) Capture-C and ChIP-seq for H3K27ac for control (grey) and after KMT2A knockdown (red) together with ChIP-seq for the N-terminus KMT2A and the C-terminus of AFF1 at the JMJD1C locus. SEM cell specific enhancer regions are highlighted in blue. Error bands represent standard error of the mean (SEM).

### Supplemental Methods

#### Cell culture

SEM cells (ACC 546) ^1^ were purchased from DSMZ (www.cell-lines.de) and cultured in Iscove’s Modified Dulbecco’s Medium (IMDM) with 10% fetal calf serum (FCS, Gibco) and Glutamax (ThermoFisher Scientific). RS4;11 (CRL-1873) cells were purchased from ATCC (www.lgcstandards-atcc.org) and cultured in RPMI 1640 with 10% FCS and Glutamax.

#### TOPmentation

10 µl protein A-coupled magnetic beads were incubated with 1 µl antibody for 4 hours at 4°C in PBS with 0.5% BSA and protease inhibitors (antibodies listed in Supplementary Table 3). Cells were fixed with 1% formaldehyde for histone modifications or double-fixed (2 mM DSG, 30 min; 1% formaldehyde, 30 min) for transcription/chromatin factors. Lysed samples (120 µl buffer with 50 mM Tris-HCl pH 8.0, 0.5% SDS, 10 mM EDTA, protease inhibitors) were sonicated to 200–300 bp fragments, neutralized with 1% Triton-X100, then pre-cleared with 5 µl protein A beads. Antibody-bound beads were washed, combined with pre-cleared chromatin, and incubated overnight at 4°C. Immunoprecipitated chromatin was washed (x3) with RIPA buffer (50 mM HEPES-KOH pH 7.6, 500 mM LiCl, 1 mM EDTA, 1% NP40 and 0.7% Na deoxycholate), then washed with Tris-EDTA and 10 mM Tris-HCl pH 8.0. Chromatin was tagmented by adding 29 µl Tagmentation Buffer (10 mM Tris-HCl pH 8.0, 5 mM MgCl2, 10% DMF) and 1 µl TDE1 (Illumina) incubating for 5 min at 37°C or 50°C (depending on the activity of the transposase lot), and stopping with 150 µl RIPA buffer. Beads were washed, and tagmented DNA amplified with NEBNext Ultra II Q5 Master Mix and indexed primers (thermal profile: 72°C 5 min, 95°C 5 min, (98°C 10 s, 63°C 30 s, 72°C 3 min) × 12, 72°C 5 min). Libraries were cleaned with Agencourt AMPure XP beads and sequenced on a NextSeq 500 or NovaSeq6000.

#### ChIP-seq/TOPmentation and ATAC sequencing analysis

Initial ChIP-seq/TOPmentation and ATAC-seq analysis was performed using the SeqNado pipeline (https://github.com/alsmith151/SeqNado). In brief, Following QC of FASTQ files by FastQC (v0.12.1;^2^), reads were trimmed using trim_galore with Cutadapt (v0.6.10;^3^). Trimmed reads were then mapped to the hg38 reference genome using Bowtie 2 (v2.5.1)^4^. PCR duplicates were removed using picard MarkDuplicates (v3.0.0; ^5^). Problematic genomic regions present in the ENCODE Blacklist ^6^ were removed from the aligned files, and further QC of the aligned files was performed using samtools (v1.17)^7^. As many of the factors that were immunoprecipitated have a mix of both sharp and broad modalities, we used the deep learning-based peak caller LanceOtron (v1.0.8)^8^ (with a peak score cut-off value of 0.5) to call peaks. Due to the limited material available, replicates were not performed and therefore an IDR-based method was not used. BigWigs were generated using the deepTools (v3.5.1) bamCoverage command (https://github.com/deeptools/deepTools), with the flags – extendReads –normalizeUsing RPKM, and visualized in the UCSC genome browser^9^. Tornado style plots were generated using deepTools (v3.5.1).

#### RNA-seq analysis

Initial RNA-seq processing was performed using the SeqNado pipline. Briefly, reads were subjected to quality checking by fastQC and trimming/QC with trim_galore. Reads were then aligned to hg38 using STAR (v2.4.2a)^10^ in paired-end mode using default parameters. Gene expression levels were quantified as read counts using the featureCounts function from the Subread package (v2.0.2)^11^ with default parameters.

#### CRISPR/Cas9-mediated enhancer deletion

Two sgRNA sequences flanking each region were designed via the integrated DNA technologies (IDT) CRISPR-Cas9 guide RNA design checker and oligos were cloned into BbsI-digested pX330-eGFP-P2A-Neo. 5×10^6^ SEM or RS4;11 cells were co-electroporated with 5 µg of each guide plasmid. After recovery for 48 hrs, transfected cells were selected with 250 µg/ml G418 (48 hrs) and then cultured on Methocult H4100 media (Stem Cell Technologies) for 14 days to isolate clones. PCR was used to screen for positive clones (Supplementary Table 4), which were confirmed by Sanger sequencing.

#### Heterogeneity in B-ALL subsets

The number of RNA-seq reads over the ncbi refseq hg38 annotation was determined using featureCounts as described above. The pyDESeq2 package ^12^ was used to perform a variance stabilising transformation and clustering/heatmap visualisation was performed using ComplexHeatmap (v 2.20.0). For ATAC-seq analysis, a consensus peak set for all samples was constructed by merging BAM files and re-calling peaks using LanceOtron to identify a set of high confidence peaks. featureCounts was used to determine the degree of accessibility at all consensus peaks for each sample. Peaks were classified into promoters (< 2.5kb from a TSS (transcription start site)) or putative enhancers (>= 2.5kb from a TSS). To compare putative enhancers and promoters, a subsample of 10, 000 elements was used to remove the class imbalance.

#### Modelling differential enhancers in SEM and RS4;11 cells

All replicates of ATAC-seq for each sample were combined and peaks were called as described. A unified set of putative enhancers and promoters were identified by intersecting ATAC-seq peaks with H3K27ac peaks and all promoter regions (peaks within 5 kb of a TSS) were filtered from the dataset. The number of reads mapping to each peak was determined using featureCounts and pyDESeq2 was used to perform the differential analysis. A putative enhancer with an FDR < 0.01 was differentially active. The read counts at all putative enhancers in the analysed ChIP-seq/TOPmentation datasets were determined using featureCounts). To cope with the imbalance in enhancer categories the imblearn (v0.8.0) package was used to randomly undersample the majority class. After class balancing a Catboost model was fitted with the catboost package (v1.2.2) and a randomised k-fold cross-validation search was used to determine the optimal parameters. The importance of each ChIP-seq dataset to the model was examined using the SHAP package (v0.39.0; ^13^).

#### Identification of ALL blast specific enhancers

To normalize immunoprecipitation efficiency across samples, scaling factors were calculated for TOPmentation data using the fit_size_factors function in pyDESeq2 on read counts from 10 kb genomic bins. BigWig files were scaled accordingly, and signal intensities were extracted over a consensus peak set for all samples. Peaks located more than 2.5 kb from a transcription start site (TSS) were classified as enhancers.

The signal from each peak was scaled to a standardized width of 1024 and inputted into an autoencoder model implemented in PyTorch (v2.1.1). The model architecture consisted of five fully connected layers, reducing the input to a 32-dimensional latent space by a factor of two with each layer, followed by a symmetric decoder. The autoencoder was trained using 41 H3K27ac bigwig/peaks sets consisting of publicly available data ^14^ and representative patient blast datasets.

To identify sample-specific enhancers, we compared peak shape and amplitude variations. Pairwise correlations of the encoded signal between samples were calculated, as well as the area under the reconstructed peak. Correlation and amplitude values were standardized, and enhancer activity was considered increased if the correlation score was less than 0 and the amplitude score exceeded 1.5. To ensure specificity, peaks were designated as sample-specific enhancers only if they showed increased activity compared to six other samples.

#### Single Cell Multiome

Single Cell multiome data was aligned using Cell Ranger ARC (v2.0). For the RNA modality, ambient RNA was removed using CellBender (v0.3.2). Samples were deconvoluted using souporcell (v2.5) and sample identity was assigned by determination of biological sex using the ratio of XIST expression compared to counts for all genes on chrY. Gene expression data was processed using scanpy (v1.10.2). Low quality cells were removed, counts were normalised, and dimensionality reduction (UMAP) was performed following the best practices workflow ^15^. Marker genes for each patient were identified using the scanpy function ‘rank_genes_groups’ using a wilcoxon test. All visualisation was performed using the scanpy package. For the ATAC-seq modality, ArchR (v1.0.2) was used for filtering low quality cells, dimensionality reduction via iterative Latent Semantic Indexing (LSI) and UMAP and peak calling. Peak calls for regions of open chromatin were generated from pseudobulk analysis of each ALL sample, followed by peak calling in MACS2. Analysis was limited to cells that were present in both RNA and ATAC modalities. Patient specific regions of open chromatin were identified using the ‘getMarkerFeatures’ function in ArchR and visualised using the ArchR package.

#### SNP calling

FastQ files were quality checked using FastQC and the reads were trimmed using Trim Galore with the parameter --2colour 20. The trimmed reads were then mapped to hg38 reference genome using Bowtie2, PCR duplicates were removed using picard MarkDuplicates (v3.0.0)^4^ and blacklisted regions^5^ were removed using bedtools. Variant calling was performed using bcftools (v1.17)^7^ the variants were split at multiallelic sites and then filtered for quality and non-heterozygous SNP variants removed. SNPs were then annotated using dbSNP (build 151)^8^.

#### Allele specific bias (ASB)

Heterozygous SNPs overlapping ATAC-seq peaks were extracted and BaalChIP (v1.26.0)^9^ was used to filter SNPs with intrinsic bias due to issues with the aligner and to identify SNPs that were over-represented for one allele.

#### Micro-Capture-C

Micro-Capture-C was performed as described 16. Briefly, 5x106 chALL1 blast cells were fixed (2% formaldehyde for 10 min), permeabilized with 0.005% digitonin (Sigma Aldrich) and snap-frozen. Thawed cells were pelleted, split into 3 aliquots and digested (typically 10-30 Kunitz U of micrococcal nuclease (NEB)) for 1 h at 37 °C before halting the reaction with 5mM EGTA. DNA was ligated in in T4 DNA ligase buffer (NEB), supplemented with 400 µM dNTPs and 5 mM EGTA in the presence of 100 U/µl DNA Polymerase I large (Klenow) fragment (NEB), 200 U/µl T4 polynucleotide kinase (NEB) and 300 U/µl T4 DNA ligase (Thermo Scientific). The reaction was incubated for 2 h at 37 °C, 8 h at 20 °C (550 rpm) before cooling to 4 °C overnight. Ligated chromatin was decrosslinked at 65oC in the presence of proteinase K (Thermo Fisher) and DNA was purified (DNeasy Blood and Tissue; Qiagen). Libraries were sonicated to a fragment size of 300 bp, and Illumina paired-end sequencing adapters (New England BioLabs, E6040, E7335 and E7500) were added. Hybridisation with oligonucleotides (Supplementary Table 5; IDT) was performed using the using the HyperCapture Target Enrichment Kit (Roche) following manufactures instructions. Analysis was performed using the MCC pipeline 17.

